# The HCN Channel Voltage Sensor Undergoes A Large Downward Motion During Hyperpolarization

**DOI:** 10.1101/522581

**Authors:** Gucan Dai, Teresa K. Aman, Frank DiMaio, William N. Zagotta

## Abstract

Voltage-gated ion channels (VGICs) underlie almost all electrical signaling in the body^1^. They change their open probability in response to changes in transmembrane voltage, allowing permeant ions to flow across the cell membrane. Ion flow through VGICs underlies numerous physiological processes in excitable cells^1^. In particular, hyperpolarization-activated cyclic nucleotide-gated (HCN) channels, which operate at the threshold of excitability, are essential for pacemaking activity, resting membrane potential, and synaptic integration^2^. VGICs contain a series of positively-charged residues that are displaced in response to changes in transmembrane voltage, resulting in a conformational change that opens the pore^3–6^. These voltage-sensing charges, which reside in the S4 transmembrane helix of the voltage-sensor domain (VSD)^3^ and within the membrane’s electric field, are thought to move towards the inside of the cell (downwards) during membrane hyperpolarization^7^. HCN channels are unique among VGICs because their open probability is increased by membrane hyperpolarization rather than depolarization^8–10^. The mechanism underlying this “reverse gating” is still unclear. Moreover, although many X-ray crystal and cryo-EM structures have been solved for the depolarized state of the VSD, including that of HCN channels^11^, no structures have been solved at hyperpolarized voltages. Here we measure the precise movement of the charged S4 helix of an HCN channel using transition metal ion fluorescence resonance energy transfer (tmFRET). We show that the S4 undergoes a significant (~10 Å) downward movement in response to membrane hyperpolarization. Furthermore, by applying constraints determined from tmFRET experiments to Rosetta modeling, we reveal that the carboxyl-terminal part of the S4 helix exhibits an unexpected tilting motion during hyperpolarization activation. These data provide a long-sought glimpse of the hyperpolarized state of a functioning VSD and also a framework for understanding the dynamics of reverse gating in HCN channels. Our methods can be broadly applied to probe short-distance rearrangements in other ion channels and membrane proteins.

---

The molecular mechanisms underlying voltage-dependent activation of VGICs remain unknown due to the lack of structural information about the hyperpolarized state of these channels^6^. HCN channels are particularly enigmatic members of the family because they are activated by membrane hyperpolarization rather than depolarization. Each HCN subunit contains an amino-terminal HCN domain (HCND), a non-domain-swapped VSD (S1-S4 helices), a pore domain (S5-S6 helices) and a carboxyl-terminal C-linker/cyclic nucleotide-binding domain (C-linker/CNBD) (Extended Data Fig. 1a)^12^. A high-resolution cryo-EM structure has been solved for the depolarized state of an HCN channel^11^, but little information exists about the conformational dynamics associated with channel activation.

To investigate VSD movement during activation of intact HCN channels in a lipid membrane, we used tmFRET to accurately measure the structure and dynamics of short-range interactions^13–16^ within a sea urchin HCN channel (spHCN)^17^. This robustly-expressing ortholog has structural and functional similarities to mammalian HCN channels^12,18^. The tmFRET measures FRET between a donor fluorophore and a non-fluorescent transition metal ion, such as Ni^2+^, Co^2+^, and Cu^2+^. Transition metal ion binding sites were introduced by site-directed mutagenesis. Efficient and specific fluorophore labeling was achieved using amber stop-codon (TAG) suppression to introduce the fluorescent, noncanonical amino acid L-Anap (Extended Data Fig. 1b)^19^. Because Anap is a small environmentally sensitive fluorophore with a short linker to the protein backbone, it is well suited as a tmFRET donor for distance measurements^15^. We also employed patch-clamp fluorometry (PCF) to allow simultaneous measurement of the structure (with tmFRET) and function (with electrophysiology) of intact spHCN channels^14^, while controlling membrane voltage and rapidly applying intracellular ligands (e.g. cAMP and transition metals) (Extended Data Fig. 1c). By combining these methods, we established a powerful way to measure the conformational dynamics associated with the gating of ion channels in their native environment.

We introduced an amber stop codon at five positions along the S4 transmembrane helix of an spHCN channel construct with a carboxyl-terminal YFP fusion (Fig. 1a and b). The five different constructs were expressed in Xenopus oocytes together with a L-Anap-specific aminoacyl tRNA synthetase/tRNA plasmid (pANAP). Fluorescence and current were recorded simultaneously using PCF (Fig. 1c and Extended Data Fig. 1c). Substantial Anap fluorescence, YFP fluorescence, and ionic current were observed from patches of all five engineered spHCN channels. Anap fluorescence was directly proportional to YFP fluorescence and ionic current (Extended Data Fig. 1c and 8). However, little or no Anap fluorescence was observed in the absence of pANAP or in the absence of the introduced amber stop codon (Fig. 1c). These results indicate that the fluorescent labeling was efficient and specific for each of the five L-Anap sites introduced into the channels.

**Figure 1.**
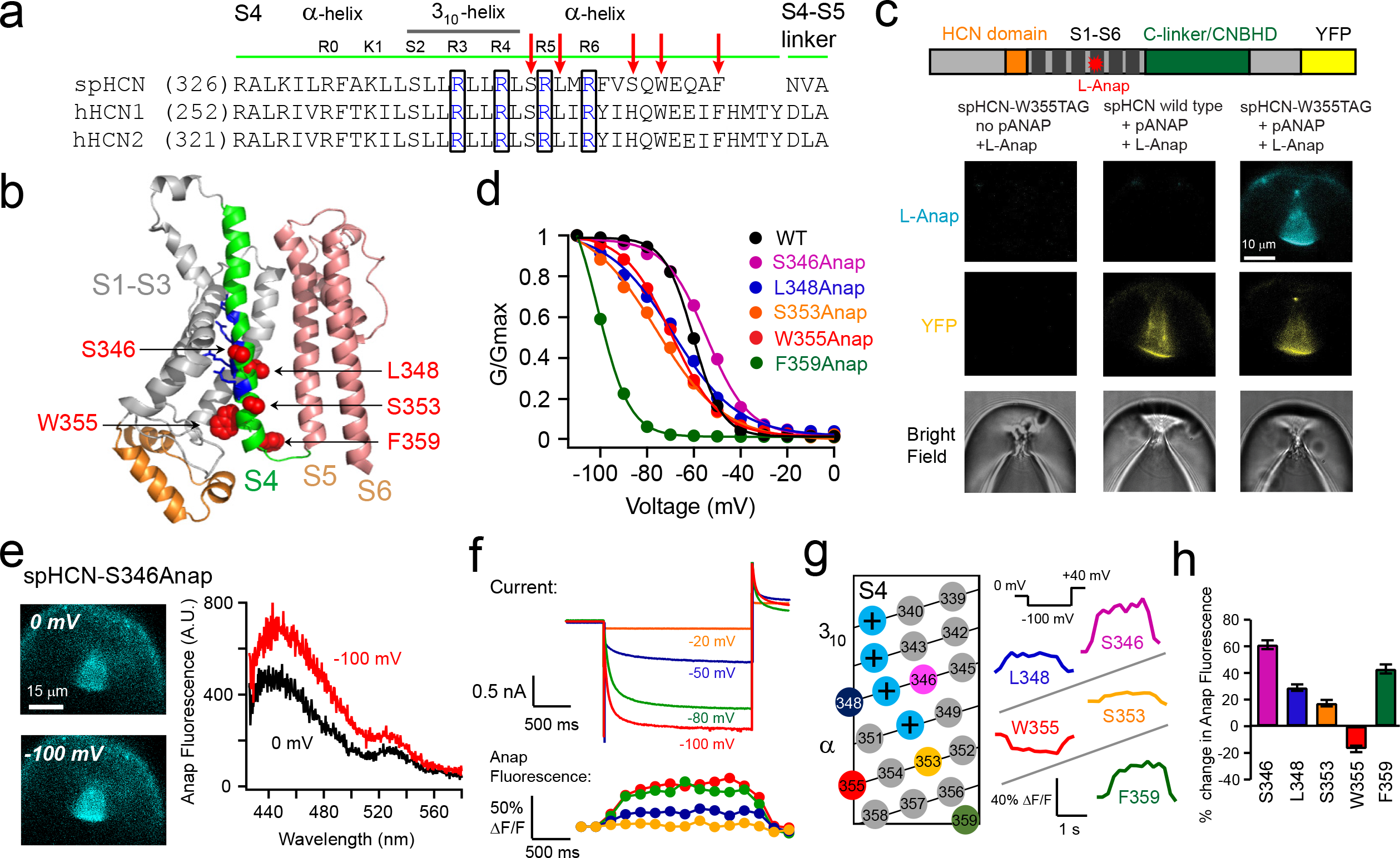
Characteristics of spHCN channels with L-Anap incorporated into the S4 voltage sensor. **a**, Sequence alignment of the S4 helix of spHCN, human HCN1 and HCN2 channels. L-Anap sites are indicated with red arrows, positively-charged voltage-sensing arginines close to the L-Anap sites are highlighted in black boxes, and the 310-helical segment of S4 is indicated with a black line. **b**, Positions of Anap sites within S4 shown in the homology model of spHCN based on human HCN1 (PDB: 5U6O). **c**, Exemplar PCF images showing the specific incorporation of L-Anap at the spHCN-W355 site. Top: Map of spHCN constructs showing eYFP fused to the C-terminal end (size of eYFP is not to scale). **d**, G-V relationships of spHCN constructs with L-Anap incorporated into different sites on S4 in the presence of 1 mM cAMP, compared to wild-type spHCN (WT). Current traces are in Extended Data Fig 2. **e**, Representative epifluorescence images and spectra of the S346Anap fluorescence at 0 mV and −100 mV. **f**, Simultaneous current and fluorescence measurements from spHCN-S346Anap channels in response to a family of hyperpolarizing voltage pulses (1 mM cAMP). **g**, Two-dimensional S4 topology and the Anap fluorescence changes at different S4 sites in response to a −100 mV hyperpolarization in the presence of 1 mM cAMP. **h**, Summary of the percent change of the Anap fluorescence for different sites within the S4 helix due to a −100 mV hyperpolarization. n = 4-7, error bars are s.e.m.

The functional properties of the engineered channels were measured using a family of hyperpolarizing voltage steps from 0 mV to −110 mV in the presence of 1 mM cAMP. All channels exhibited appreciable hyperpolarization activation (Extended Data Fig. 2). The conductance-voltage (G-V) relationships were similar to that of wild-type spHCN channels in four of the channels (Fig. 1d). However, the F359Anap site at the end of S4 resulted in activation kinetics that were substantially slower, and a G-V curve that was shifted by more than 30 mV in the hyperpolarizing direction, in comparison to wild-type spHCN (Fig. 1d, Extended Data Fig. 2). These results suggest that, with the exception of F359Anap, introduction of L-Anap into the S4 helix did not substantially perturb the hyperpolarization-dependent gating of these channels.

The environmental sensitivity of Anap fluorescence allowed us to test for any changes to the environment surrounding the S4 helix during voltage-dependent activation (hyperpolarization) of spHCN channels. For spHCN-S346Anap, there was a substantial increase in fluorescence during hyperpolarizing voltage pulses to −100 mV (61.2 ± 3.3%, n = 6) (Fig. 1e,f and Supplementary Video 1). This increase was accompanied by a shift in the peak emission spectrum to longer wavelengths (8.3 ± 1.5 nm) (Extended Data Fig. 7a,f), suggesting that S346Anap enters a more hydrophilic environment at −100 mV. An increase in fluorescence was also observed for L348Anap, S353Anap, and F359Anap, and a decrease observed for W355Anap (Supplementary Video 2), with or without a shift in the peak emission (Fig. 1g,h and Extended Data Fig. 7). For S346Anap, the time course and voltage dependence of these fluorescence increases matched the time course and voltage-dependence of the ionic currents (Fig. 1f,g and Extended Data Fig. 3). In contrast, an Anap site in the S1 transmembrane segment (W218Anap) exhibited little or no fluorescence change when hyperpolarized to −100 mV (Extended Data Fig. 4b and Extended Data Fig. 7e). These results suggest that the fluorescence changes at the S4 Anap sites are reporting, at least in part, the rearrangement of the VSD during channel activation.

In order to directly measure the direction of S4 movement, we measured Anap fluorescence in the presence of the transition metal Co^2+^, whose absorption spectrum overlaps with the emission spectra of L-Anap (Fig. 2a)^13^. A transition metal ion binding site was introduced into the spHCN-S346Anap channel as a dihistidine (diHis) motif (L182H, L186H) on an α helix in the HCN domain, directly below S4 (Fig. 2b). Upon application of 1 mM Co^2+^, there was substantial voltage-dependent quenching of Anap fluorescence (Fig. 2c). The quenching was reversible by EDTA application and was substantially less in spHCN-S346Anap channels lacking the diHis site, indicating that it arose from FRET between S346Anap and Co^2+^ bound to the diHis site in the HCN domain. Importantly, Co^2+^-dependent quenching was much greater at hyperpolarized voltages than at depolarized voltages (Fig. 2c). We quantified the apparent FRET efficiency by calculating the fractional decrease in Anap fluorescence in 1 mM Co^2+^ and correcting for the small quenching in spHCN-S346Anap lacking the diHis site. The FRET efficiency increased by ten-fold, from 0.03 ± 0.01 at 0 mV to 0.30 ± 0.01 at −120 mV (Fig. 2d). These results suggest that the S346 position in the S4 helix moves downward – closer to the HCN domain – during hyperpolarization activation of HCN channels.

**Figure 2.**
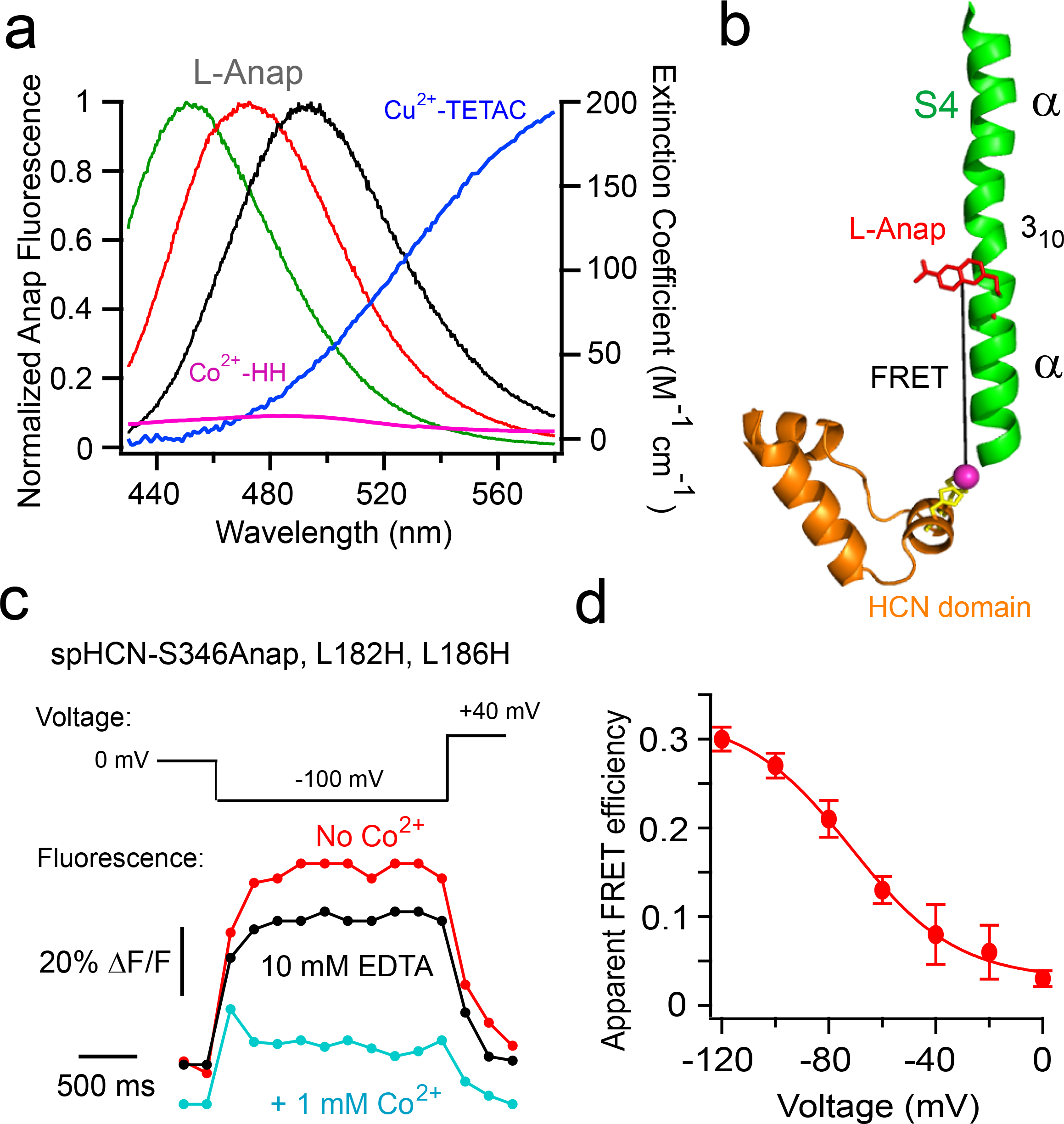
tmFRET detects a hyperpolarization-dependent downward movement of S4 in spHCN-346Anap channels. **a**, Spectral properties of free L-Anap emission and transition metal ion absorption. Emission spectra from L-Anap in different solvents (black: SBT buffer, red: EtOH, green: DMSO) are overlaid on the absorption spectra of Cu^2+^-TETAC (dark blue line) and Co^2+^-HH (magenta line) **b**, Cartoon showing FRET between the S346Anap site and the diHis site in the HCN domain (L182H-L186H). **c**, Time course of Anap fluorescence in spHCN-S346Anap, L182H, L186H channels in the presence of cAMP before and after Co^2+^ application and EDTA to sequester Co^2+^. **d**, Voltage dependence of the apparent FRET efficiency for the spHCN-S346Anap, HH construct in the presence of 1 mM cAMP (n = 3).

We switched to a new method for tmFRET, called ACCuRET (Anap Cyclen-Cu^2+^ resonance energy transfer), to measure the movement of the S4 helix more precisely. Instead of a diHis motif, ACCuRET introduces a higher affinity transition metal ion-binding site using TETAC (1-(2-pyridin-2-yldisulfanyl)ethyl)-1,4,7,10-tetraazacyclododecane) (Fig. 2a and Fig. 3a)^20^. TETAC is a cysteine-reactive compound with a short linker to a cyclen ring that binds transition metal ions with subnanomolar affinity^21^. Single cysteines introduced into the protein react with TETAC to create a high-affinity metal binding site for tmFRET. This binding site has two major advantages over diHis: 1) it avoids using high (millimolar) concentrations of transition metals, which could bind to other sites or have unwanted effects on the protein; 2) TETAC increases and shifts the absorbance of Cu^2+^ to extend the range of distances that can be measured with tmFRET^20^ (Fig. 2a). These advantages recently enabled the accurate measurement of absolute distances and distance changes in protein backbones that are associated with structural rearrangements in soluble and membrane proteins^20^.

**Figure 3.**
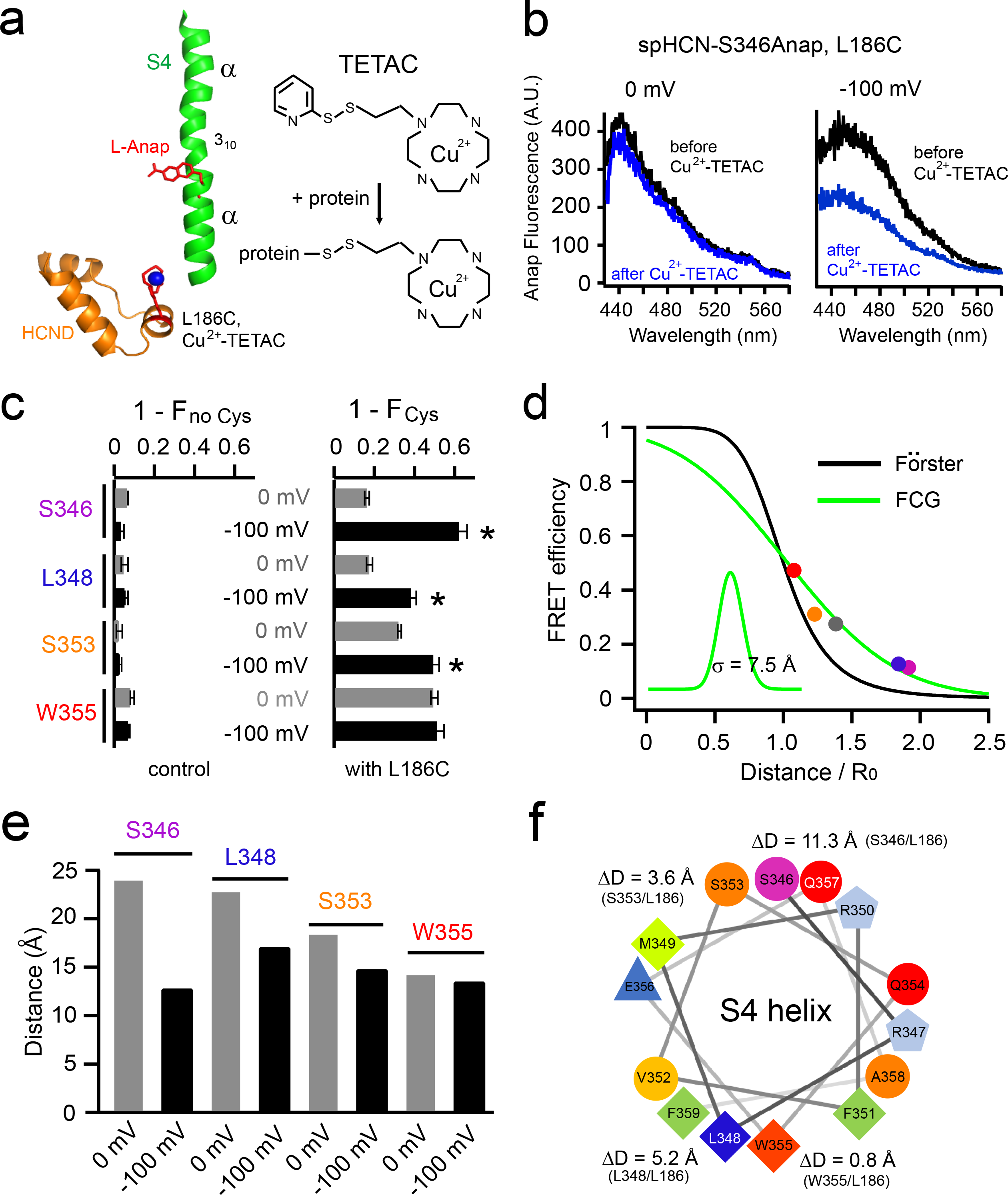
The distance change measured by ACCuRET decreases as Anap is positioned closer to the C-terminal end of S4. **a**, Cartoon showing ACCuRET between S346Anap and Cu^2+^-TETAC attached to L186C in the HCN domain. **b**, Representative spectra of the Anap emission at 0 mV and −100 mV, before and after quenching by Cu^2+^-TETAC, for spHCN-S346Anap, L186C channels. **c**, Summary of the fraction of Anap fluorescence quenched by Cu^2+^-TETAC for the 4 Anap sites in S4, without (left, n = 2-4) and with (right, n = 4-7) the introduced cysteine L186C. **d**, Distance dependence of the measured FRET efficiency at 0 mV for four L-Anap sites in S4 and the W218Anap site in S1 versus the β-carbon distances between each site and L186C (based on the spHCN homology model). Also shown are the predicted distance dependencies of the Förster equation (black) and the Förster Convolved with Gaussian (FCG) equation (green), using a Gaussian distribution with a standard deviation σ of 7.5 Å (inset). **e**, Summary of the distances of each FRET pair at 0 mV and −100 mV, calculated from the FCG plot in panel d, using the FRET efficiencies in Extended Data Fig 8b. **f**. Helical-wheel display of the C-terminal α-helical part of S4 (S346 to F359) viewed from the extracellular side, highlighting the magnitude of the hyperpolarization-induced distance changes for the respective FRET pairs in panel e.

We employed ACCuRET to measure the size of the movement of the S4 helix during hyperpolarization activation of spHCN. In combination with the Anap sites, we introduced a single cysteine (L186C) into the HCN domain of cysteine-depleted spHCN constructs (see Methods). Neither mutating the unwanted cysteines nor Cu^2+^-TETAC labeled L186C had an appreciable effect on hyperpolarization-dependent gating (data not shown). Similar to the effect of Co^2+^ on the diHis motif, the addition of Cu^2+^-TETAC caused a significant reduction in Anap fluorescence for spHCN-S346Anap, L186C, particularly at −100 mV (Fig. 3b, c). This Cu^2+^-TETAC quenching had no effect on the shape of the Anap emission spectrum and was negligible in the absence of the introduced cysteine, indicating that it arose from FRET between S346Anap and Cu^2+^-TETAC bound to L186C on the HCN domain (Fig. 3b, c). Like the effect of Co^2+^ on the diHis motif, the quenching was much greater at −100 mV (62.5 ± 3.6 %) than at 0 mV (16.7 ± 0.8 %) (Fig. 3c). Unexpectedly, this voltage-dependent quenching substantially decreased as Anap sites became closer to the C-terminal end of S4 helix (Fig. 3c). These data imply that S4 is not simply translating as a rigid body directly towards the HCN domain during hyperpolarization activation.

To investigate the nature of this S4 motion in more detail, we used FRET efficiencies to calculate the distances between each Anap site and L186 at 0 mV and −100 mV. FRET efficiencies were calculated from the fractional decrease in Anap fluorescence after application of Cu^2+^-TETAC and corrected for the small Cu^2+^-TETAC-dependent quenching in Anap constructs lacking L186C (Extended Data Fig. 6). We also determined R0 values (the distance predicted to produce 50% FRET efficiency) for each site at both voltages, using emission spectra and fluorescence intensity data (Extended Data Fig. 7 and 8). The measured FRET efficiencies for each site at 0 mV were plotted against the β-carbon distances between each FRET pair (in distance/R0 units) measured from an spHCN homology model built from the cryo-EM structure of hHCN1 (also at 0 mV) (Fig. 3d). The FRET efficiencies decreased with distance, but as previously observed for tmFRET^20^, the distance dependence was shallower than predicted by the Förster equation. This shallower distance dependence has been attributed, in part, to heterogeneity of the interatomic distances in proteins. Förster Convolved Gaussian (FCG) theory accounted for this heterogeneity and provided an empirical measurement of the distance dependence in our experiments (Fig. 3d).

Using our calibrated distance dependence, FRET efficiencies, and R_0_ values, we calculated the distances for each FRET pair at each voltage (Fig. 3e). As expected, the distances at 0 mV closely matched the distances in the homology model based on the hHCN1 structure at 0 mV (RMSD = 1.0 Å). However, the distances at −100 mV deviated significantly from the homology model. The change in distance decreased steadily along the S4 helix from 11.3 Å at S346 to only 0.8 Å at W355. These results support our proposal that the movement of S4 is not a rigid body movement directly towards L186 in the HCN domain. Furthermore, since both S346 and S353, and both L348 and W355, are on the same side of the α helix (Fig. 3f), no combination of rotation and translation towards L186 can explain our measured distance changes. Instead, our results suggest that the S4 helix moves downward and tilts during hyperpolarization activation.

The interpretation above assumes that L186 on the HCN domain does not move substantially during activation. To test for movement of the HCN domain, we measured tmFRET between L186C and W218Anap on the S1 helix, and calculated distances at 0 mV and −100 mV as above (Extended Data Fig. 4d,e). Interestingly, W218Anap moved a little closer to the HCN domain at −100 mV (ΔD = 2.7 Å). This could reflect a small movement of the HCN domain, or a subtle downward movement of S1, or a combination of the two. In any case, these small movements were included in our structural modeling below, and did not appreciably affect our measurements of the much larger movements of some sites on the S4 segment.

To estimate the actual movement of the S4 helix, we used our distances as constraints in Rosetta-based structural modeling of the conformational change of the VSD (Fig. 4a). The models at 0 mV and −100 mV accurately reproduced the distances and distance changes measured from our ACCuRET experiments (Fig. 4b). At 0 mV, the model was very similar to the spHCN homology model based on the HCN1 cryo-EM structure at 0 mV (Supplementary Video 4). However, with the −100 mV constraints, the model revealed a large downward movement of S4 of about two turns of the helix (Fig. 4a and Supplementary Video 3). In addition, the C-terminal end of the S4 helix tilted away from the HCN domain, producing a kink in the helix at position R344 (Fig 4a). Little or no rotation of S4 was observed in the model (indeed, rotation caused a poorer fit to our measured distances), and the HCN domain underwent only a small upward movement. Furthermore, extensive salt bridges between S4 arginines and S1-S3 aspartic acids were seen at both voltages (Fig. 4c). The 3_10_-helical nature of S4 in the gating charge transfer center (indicated in Fig. 1a)^22^ positions the voltage-sensing arginines R341 and R344 in line on the same side of S4 at both voltages. Therefore, this region moves almost vertically to exchange their ion-pair partners without much rotation (Fig 4c and Supplementary Video 5). Together, our experiments provide an explicit model of the conformational dynamics of the spHCN VSD during hyperpolarization activation, based on experimentally-determined distance measurements under native conditions.

**Figure 4.**
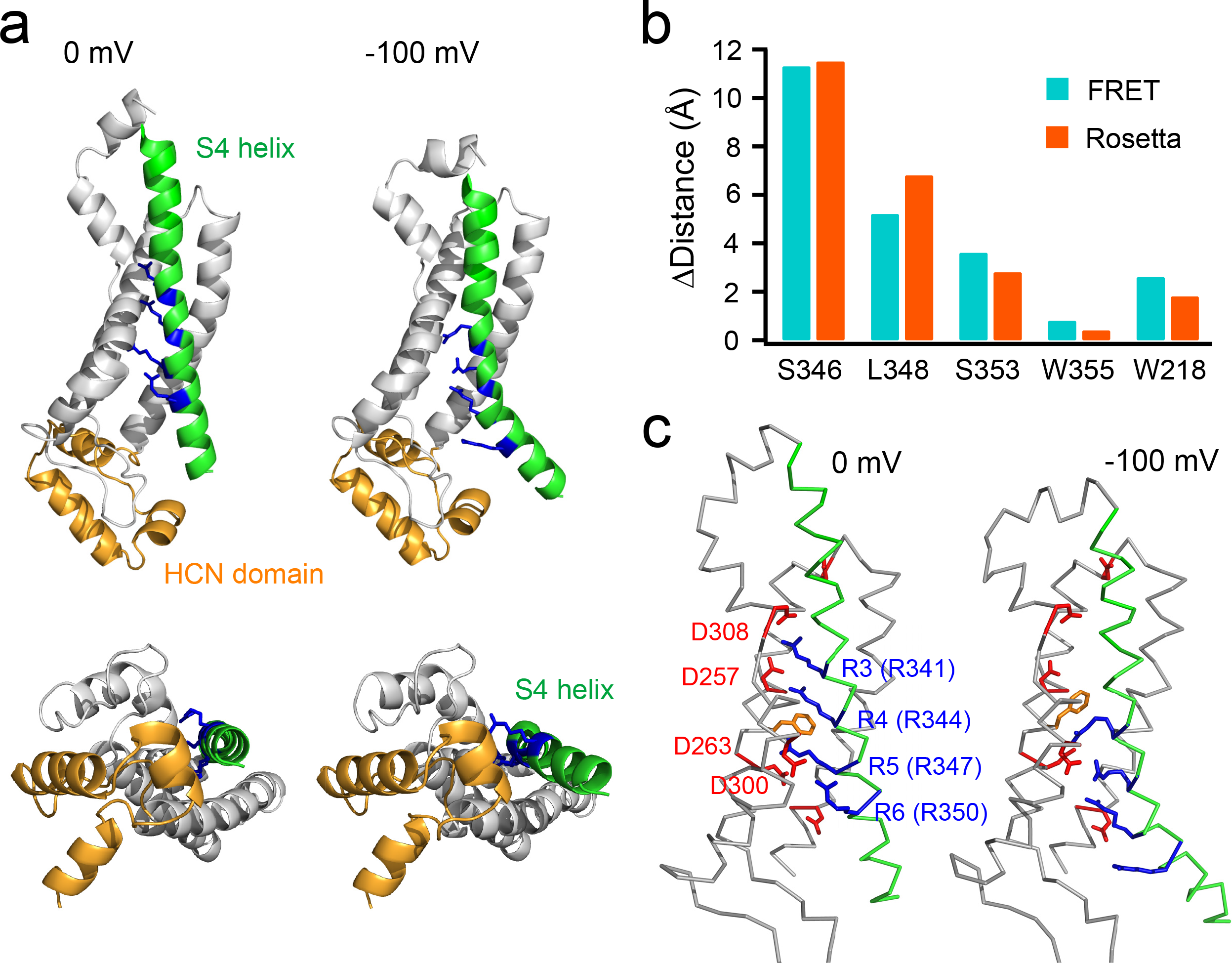
Rosetta-based model of S4 movement in HCN channels. **a**, Model structures of the voltage-sensor domain of spHCN at rest (0 mV) and hyperpolarized (−100 mV) voltages. Top: side view parallel to the membrane. Bottom: view from the intracellular side. The S1-S3 helices are in a surface presentation. **b**. Comparison of the measured distance changes using tmFRET from Fig. 3 and the distance changes in the Rosetta models. **c**. Structural diagrams showing the ion pair partners between arginines (blue) within S4 and aspartic acids (red) within S1-S3 at 0 mV and −100 mV in the Rosetta model. D257 and D308 interact with R341 and R344 within the charge transfer center at 0 mV whereas D263 and D300 interact with these two arginines at −100 mV. The “phenylalanine cap” F260 in the hydrophobic constriction site of the charge transfer center is highlighted in orange.

This new model of the movement of the S4 helix could further unify the mechanisms of VSD activation for both hyperpolarization-and depolarization-activated channels^6,7^. A downward movement corresponding to two turns of the S4 helix could produce an equivalent charge movement of up to two electronic charges per subunit (eight charges per channel) during voltage-dependent activation of spHCN. This is more than enough charge movement to reproduce the steepness of the G-V, Q-V, and F-V curves for these channels^7,23^. The tilting motion of S4 is also consistent with previous cysteine accessibility experiments in HCN channels^24,25^. Furthermore, the rearrangement is similar to the S4 movement in VSD2 of the depolarization-activated two-pore channel 1 (TPC1) inferred from structures of mouse TPC1 (up state) and Ba^2+^-inhibited Arabidopsis TPC1 (down state) at 0 mV^26,27^ (Supplementary Video 6), but is incompatible with other models proposing that the S4 helix is relatively immobile^28^. Moreover, considering the tight packing of S4 and S5 helices in HCN channels^11^, the downward and tilting movement of the S4 helix could bypass the short S4-S5 linker to influence S5 directly, thus allowing the channels to open^29,30^. Our study has revealed, for the first time, the detailed movement of the S4 helix of a VSD during hyperpolarization. It also establishes a methodological framework to decipher the reverse gating mechanism of HCN channels.

## METHODS

### Molecular Biology

The full-length spHCN cDNA (a gift from U. B. Kaupp, Molecular Sensory Systems, Center of Advanced European Studies and Research, Bonn, Germany; GenBank: Y16880) was subcloned into a modified pcDNA3 vector (Invitrogen, Carlsbad, CA) that contained a carboxy-terminal eYFP, a T7 promoter and 5’ and 3’ untranslated regions of a *Xenopus* β-globin gene. For the ACCuRET experiments, a cysteine-depleted spHCN construct was used with the following mutations: C211A, C224A, C369A, C373A. Point mutations were made using Quickchange II XL Site-Directed Mutagenesis kit (Agilent technologies, Santa Clara, CA). The sequences of the DNA constructs were confirmed by fluorescence-based DNA sequencing (Genewiz LLC, Seattle, WA). The RNA was synthesized in vitro using HiScribe T7 ARCA mRNA Kit (New England Biolabs, Ipswich, MA) or mMESSAGE mMACHINE T7 ULTRA Transcription Kit (ThermoFisher, Waltham, MA) from the linearized plasmid.

### Oocyte Expression and Electrophysiology

The animal-use protocols were consistent with the recommendations of the American Veterinary Medical Association and were approved by the Institutional Animal Care and Use Committee (IACUC) of the University of Washington. *Xenopus* oocytes were prepared as previously described^31^. L-Anap (free-acid form, AsisChem, Waltham, MA) was made as a 1 mM stock in water at a high pH by adding NaOH and stored at −20°C. The pANAP plasmid (purchased from Addgene, Cambridge, MA) contained the orthogonal tRNA/aminoacyl-tRNA synthetase specific to L-Anap^19^. The pANAP plasmid (~40 nL of 200 ng/μl) was injected into the *Xenopus* oocyte nucleus. Then, L-Anap (~50 nL of 1 mM) and channel mRNA were injected separately into the oocyte cytosol.

Channel currents were recorded from the oocytes 2 to 4 days after injection using the inside-out configuration of the patch-clamp technique with an EPC-10 (HEKA Elektronik, Germany) or Axopatch 200B (Axon Instruments, Union City, CA) patch-clamp amplifier and PATCHMASTER software (HEKA Elektronik). Inside-out patch clamp recordings were made approximately 5 mins after patch excision, when the run-down of HCN channels due to PI(4,5)P_2_ depletion was almost complete^32^. Borosilicate patch electrodes were made using a P97 micropipette puller (Sutter Instrument, Novato, CA). The initial pipette resistances were 0.3–0.7 MΩ. Recordings were made at 22°C to 24°C. A μFlow microvolume perfusion system (ALA Scientific Instruments, Farmingdale, NY) was used to change solutions during the experiment. All recordings were made in the presence of 1 mM cAMP on the cytosolic side of the patch. For oocyte patch-clamp recording and Co^2+^-HH tmFRET experiments, the standard bath and pipette saline solutions contained 130 mM KCl, 10 mM HEPES, 0.2 mM EDTA, pH 7.2. 1 mM CoSO_4_ was added to the perfusion solution with EDTA eliminated. For ACCuRET experiments, stabilization buffer (SBT) (130 mM KCl; 30 mM Trizma Base; 0.2 mM EDTA, pH 7.4) was used for the bath solution. The Cu^2+^-TETAC was prepared as previously described^20^. TETAC (Toronto Research Chemicals, Toronto, Canada) was made as a 100 mM stock in DMSO and stored at −20°C. On the day of the experiment, 1 μL each of 100 mM TETAC stock and 110 mM CuSO4 stock were mixed together and incubated for one minute until the solution turned to a deeper shade of blue, indicating binding of Cu^2+^ to the TETAC. To this mixture, 98 μL of SBT was added, giving a solution of 1.1 mM Cu^2+^ and 1 mM TETAC. The 10% over-abundance of Cu^2+^ ensured that all the TETAC was bound with Cu^2+^. This solution was then diluted 1:100 in SBT buffer, giving a final concentration of 10 μM Cu^2+^-TETAC, and perfused on to the patch.

### Fluorescence Measurements

Patch-clamp fluorometry (PCF) simultaneously records fluorescence and current signals in inside-out patches. Our PCF experiments were performed using a Nikon Eclipse TE2000-E inverted microscope with a 60X 1.2 NA water immersion objective. Epifluorescence recording of L-Anap was performed with wide-field excitation using a Lambda LS Xenon Arc lamp (Sutter Instruments) and a filter cube containing a 376/30 nm excitation filter and a 460/50 nm emission filter. A 425 nm long-pass emission filter was used for the spectral measurement of L-Anap. YFP was measured with a filter cube containing a 490/10 nm excitation filter and a 535/30 nm emission filter. Images were collected with a 200 ms exposure using an Evolve 512 EMCCD camera (Photometrics, Tucson, AZ) and MetaMorph software (Molecular Devices, Sunnyvale, CA). The Anap fluorescence at the −100 mV steady state for the ACCuRET experiments were captured a few seconds after switching the voltage from 0 mV to −100 mV. For a comparison of the Anap and YFP fluorescence intensities, the settings of the EMCCD camera were kept the same.

For spectral measurements, images were collected by a spectrograph (Model: 2150i, 300 g/mm grating, blaze = 500 nm; Acton research, Acton, MA) mounted between the output port of the microscope and the Evolve 512 EMCCD camera. The membrane patch was positioned in the entrance slit of the spectrograph with the pipette parallel to the slit. The slit width was made slightly smaller than the width of the patch to block light from regions of the patch attached to the glass. The slit width we used had little or no effect on the shape of the emission spectrum (Extended Data Fig. 5b). The spectrograph produced an image on the camera where the vertical dimension was positioned along the axis of the pipette, and the horizontal dimension was wavelength. The wavelength scale was calibrated with known filters and laser lines. Anap spectra were recorded by measuring line-scans through the patch area, background subtracted using line scans through the non-fluorescent region in the pipette outside of the patch, and corrected for the filter characteristics of the spectrograph.

Fluorometry experiments on free L-Anap in solvents were performed using a Jobin Yvon Horiba FluoroMax-3 spectrofluorometer (Edison, NJ). Absorption spectra were recorded with a DU 800 spectrophotometer (Beckman Coulter, Brea, CA). Sub-micro fluorometer cuvettes (100 μL Starna, Atascadero, CA) were used for both fluorometry and spectrophotometry. For emission spectra of free L-Anap-ME (AsisChem), an excitation wavelength of 350 nm and 1 nm slits for excitation and emission were used.

### Rosetta Modeling

A homology model (State 0) of spHCN channel based on the HCN1 structure^11^ (PDB: 5U6O) was created using RosettaCM without any experimental constraints^33,34^. To obtain a model of the spHCN channel at 0 mV and −100 mV, models were refined using RosettaCM with three types of constraints applied to the homology model: (1) Coordinate constraints (tethering the Cartesian coordinates of an atom) were applied on every α-carbon atom except for those in the S4 helix. These were “top-out” constraints, which are roughly harmonic up to 1 Å from the minimum, and flat beyond; (2) harmonic pairwise distance constraints were applied on all backbone hydrogen bonding groups; and (3) experimentally-derived distances at 0 mV (State 1) and −100 mV (State 2) based on our tmFRET measurements were applied as “flat-bottom” harmonic constraints. For setting the constraint weights for a satisfactory model, we performed a grid search of the weights on each of the three constraint types. The weakest overall constraints that led to satisfied experimental constraints in both states were used: a weight of 15 on the experimental data, and a weight of 2 on the hydrogen-bond and coordinate constraints. All modelling was carried out in the context of the C4 tetramer. Structural representations and morphs were created using the PyMOL software (https://pymol.org). A structural morph was created between the homology model and State 2 (Fig 4a and Supplementary Video 3), highlighting the hyperpolarization-dependent movement of S4. A structural morph between the homology model and State 1, based on the FRET distance measurement at 0 mV, displays very little movement (morph in Supplementary Video 4), indicating the distance measurement from tmFRET is compatible with the homology model and the structure of HCN1.

### Data Analysis

Data were analyzed using IgorPro (Wavemetrics) and Image J (NIH). Data parameters were expressed as mean ± s.e.m. of n experiments from distinct samples. Statistical significance (\**p* < 0.05) was determined by using Student’s *t* test.

The conductance-voltage (G-V) relationships were measured from the instantaneous tail currents at +40 mV following voltage steps from 0 mV to between 0 and −110 mV. The leak tail currents following steps to +40 mV were subtracted, and the currents were normalized to the maximum tail current following voltage steps to −110 mV (G/G_max_). The relative conductances were plotted as a function of the voltage of the main pulse and fitted with a Boltzmann equation:

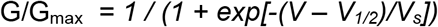

where *V* is the membrane potential, *V*_*1/2*_ is the potential for half-maximal activation, and *V*_*s*_ is the slope factor.

Fluorescence images from the membrane patches were imported into ImageJ^35^ for analysis. Regions of interest were drawn by hand around the dome of the patch, excluding any regions that appeared attached to the glass. For each patch, a background region was selected in the pipette outside of the patch. The mean gray value of the background region of interest was subtracted from the mean gray value of the region of interest of the patch to give the mean fluorescence intensity (*fl*).

The FRET efficiency was calculated using the following equations (E1) as previously described^36^:

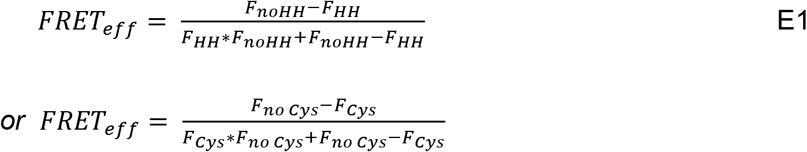

Where *F*_*HH*_ and *F*_*no HH*_ are the fractions of fluorescence that are unquenched by Co^2+^ in channels with and without HH sites, respectively, e.g. 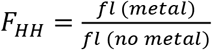. Similarly, *F*_*Cys*_ and *F*_*no Cys*_ are the fractions of fluorescence that are unquenched by Cu^2+^-TETAC in protein with and without cysteines respectively. These equations for calculating FRET efficiency assume that the decrease in *F*_*no HH*_ is due to nonspecific energy transfer such as solution quenching or FRET to a metal ion bound to an endogenous metal binding site.

Alternatively, the FRET efficiency was calculated using the following equations (E2), which only account for nonspecific decreases in fluorescence (e.g. photobleaching) but not energy transfer.

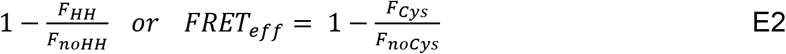

The FRET efficiencies produced by E1 and E2 were similar because of the low degree of background quenching^20,36^ (Extended Data Fig. 6). To calculate the mean and standard error of the mean for the FRET efficiency, we used the mean and standard error of the mean for our fractional quenching measurements (i.e. *F*_*HH*_, *F*_*no HH*_, *F*_*Cys*_, and *F*_*no Cys*_) in Monte Carlo resampling (1 × 10^6^ cycles; NIST Uncertainty Machine v1.3.4; https://uncertainty.nist.gov).

The R_0_ values, the distances that predict 50% energy transfer, were calculated using the established equation as previously published^20^.

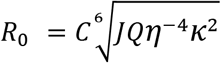

Where *C* is the scaling factor, the index of refraction *η* was 1.33 and the orientation factor *κ*^2^ was assumed to be 2/3, a reasonable assumption for an isotropic metal ion^37^. The R_0_ for each L-Anap site paired with Cu^2+^-TETAC were calculated using the overlap integral *J* of the donor emission spectrum (Extended Data Fig 7) and the absorption spectrum of Cu^2+^-TETAC, as well as the estimated quantum yield of L-Anap at that site (at 0 mV or −100 mV) (Extended Data Fig 8). Since the shape of the Anap emission spectrum does not change appreciably in different solvents (Extended Data Fig. 5a inset), we shifted the free L-Anap spectrum measured in the cuvette to match the peak of the emission spectra measured on the microscope for each Anap site at 0 mV and −100 mV (Extended Data Fig 7). This strategy removes the contamination of the Anap emission spectrum by the YFP emission peaked at around 530 nm. The YFP emission is most likely due to the direct excitation of YFP by the Anap excitation, although the possibility of a small amount of FRET between Anap and YFP cannot be completely excluded. Finally, the relative quantum yields of L-Anap at each site at 0 mV and −100 mV were estimated from the slopes of the plots of Anap intensity vs. YFP intensity at 0 mV (Extended Data Fig 8) and the relative brightness at −100 mV vs. 0 mV (*fl*_*Anap, −100 mV*_ / *fl*_*Anap, 0 mV*_) (Fig. 1h). For each site, Quantum yield (0 mV) = 0.3 * Slope (Anap vs YFP) (Extended Data Fig 8 a-e) and Quantum yield (−100 mV) = Quantum yield (0 mV) * (*fl*_*Anap, −100 mV*_ / *fl*_*Anap, 0 mV*_). This estimate of quantum yield assumes that the change in brightness of Anap is primarily due to a change in quantum yield and not extinction coefficient, as appears to be true for the solvents that were tested (Extended Data Fig. 5c). In addition, the quantum yields estimated fell within the range of the quantum yield of L-Anap in different solvents (0.2-0.5)^15,20^. R0 values calculated using these estimated spectral overlaps and quantum yields were as follows: 13.1 Å (0 mV) and 14.9 Å (−100 mV) for S346Anap; 12.6 Å (0 mV) and 13.5 Å (−100 mV) for L348Anap; 13.8 Å (0 mV) and 14.1 Å (−100 mV) for S353Anap; 13.3 Å (0 mV) and 13.0 Å (−100 mV) for W355Anap; 13.9 Å (0 mV) and 13.9 Å (−100 mV) for W218Anap.

The distances between L-Anap and the metal ion were determined using the FRET efficiencies calculated with E1 and the R_0_ values calculated above. The empirical distance dependence of FRET was determined by plotting the measured FRET efficiencies at 0 mV for each site as a function of the beta carbon distances (in R_0_ units) determined from the homology model based on the cryo-EM structure of HCN1. These data were then fit with a Förster equation convolved with a Gaussian distribution (FCG) to account for heterogeneity in the distances between the donor and acceptor and the shallower distance dependence, as previously described^20^. The wide Gaussian distribution used for the FCG (standard deviation σ = 7.5 Å) is possibly due, in part, to the flexibility of the short helix in the HCN domain where L186 is located. This FCG function provided an empirical ruler for estimating the distances for each of the Anap sites at 0 mV and −100 mV.

## Data availability

Data are available from the corresponding author upon reasonable request.

## Acknowledgments

We thank Ximena Optiz-Araya and Sarie Haraguchi for animal care and surgical support; Sharona E. Gordon, Lucie Delemotte, and all members of the Zagotta laboratory for their helpful advice and support, and Lesley Anson for comments on the manuscript. This work was funded by NIH Grants R01EY010329, R01MH102378, R01GM125351, and American Heart Association Award 14CSA20380095 (to W.N.Z.) and NIH Grant F32NS077622 (to T. K. A.).

## Author Contributions

G.D., T.K.A, and W.N.Z. conceived and designed experiments, G.D. performed experiments. T.K.A. performed pilot experiments. F.D. designed and performed Rosetta-based computational modeling. G.D., T.K.A., F.D., and W.N.Z. analyzed data. W.N.Z and G.D. wrote the manuscript. G.D., T.K.A, F.D., and W.N.Z. edited the manuscript.

---

**Extended Data Fig. 1.**
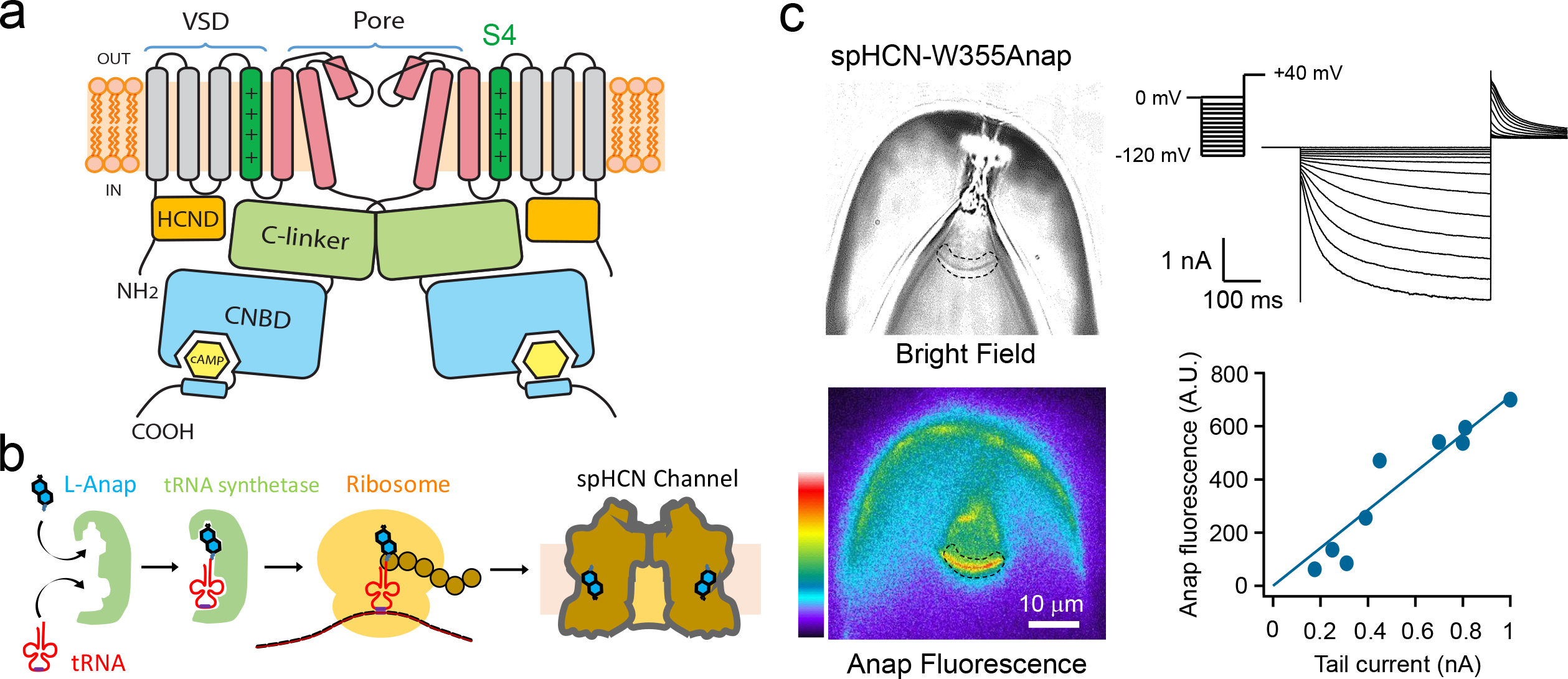
Patch-clamp fluorometry (PCF) simultaneously measures current and Anap fluorescence in HCN channels. **a**, Cartoon showing the architecture of an HCN channel subunit (only two subunits are shown). Each subunit contains a cyclic nucleotide-binding domain (CNBD) in the carboxy-terminal region that is connected to the pore region via a gating ring formed by the C-linker of each subunit. L-Anap is incorporated into the S4 helix in the voltage sensor domain. **b**, Cartoon illustrating amber stop-codon suppression strategy for incorporating noncanonical amino acids. **c**, Representative PCF images showing a giant inside-out patch expressing spHCN-W355Anap channels in bright field (top) and a heat-map presentation of the fluorescence from the same patch (bottom), as well as the channel current elicited by a series of hyperpolarizing voltages from 0 mV to −120 mV. The dashed area is an exemplar region of interest used for quantifying the patch fluorescence. The intensity of Anap fluorescence is proportional to the magnitude of the peak tail current.

**Extended Data Fig. 2.**
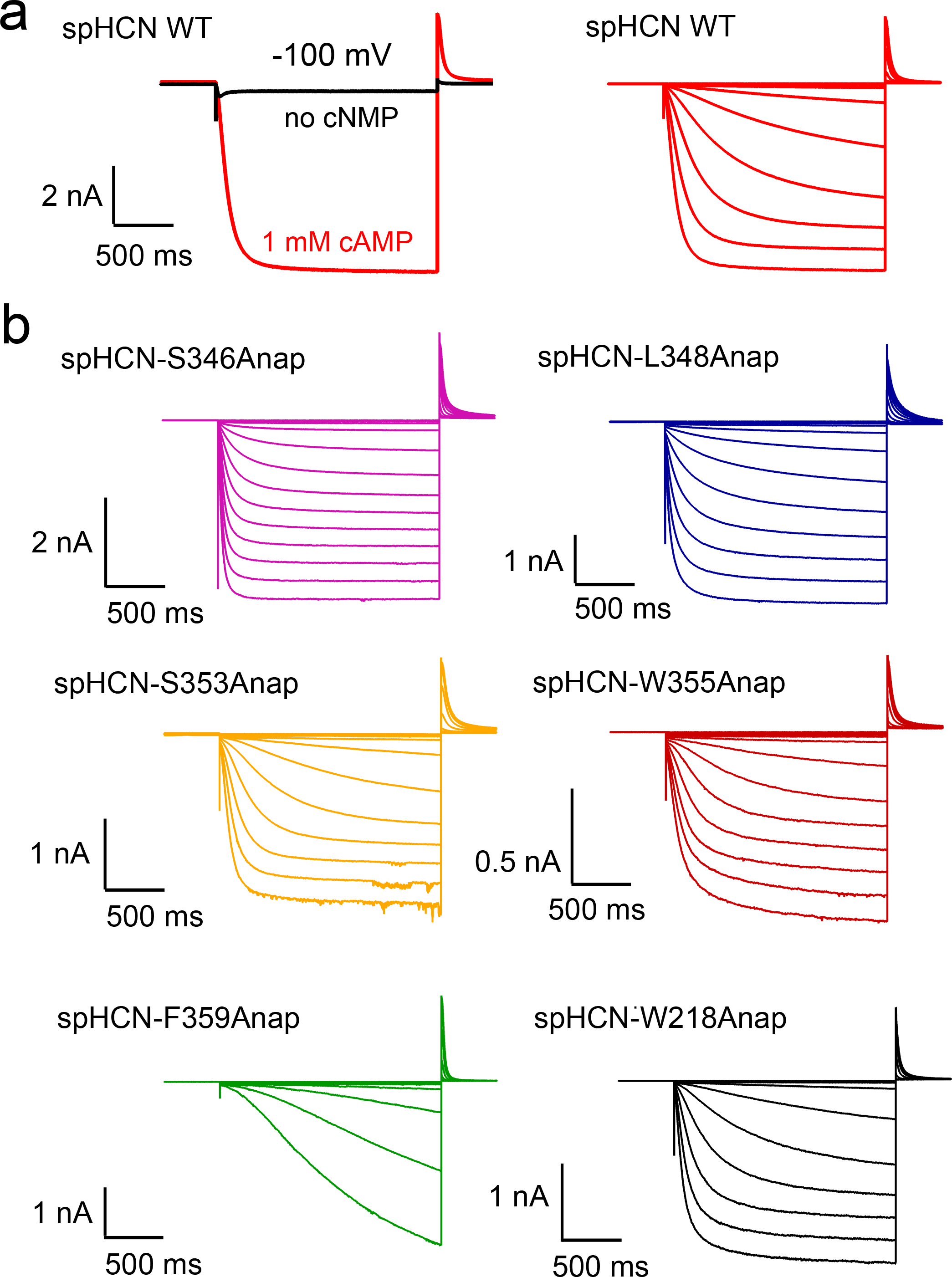
Electrophysiological properties of spHCN channels with incorporated Anap. **a**, Left: representative current traces of wild-type spHCN channels in the absence or the presence of 1 mM cAMP. In the absence of cyclic nucleotide, the spHCN channel rapidly inactivates with hyperpolarization. Right: representative current traces from wild-type spHCN channels elicited by a series of hyperpolarizing voltage pulses from 0 mV to −120 mV, in the presence of 1 mM cAMP. **b**, Representative current traces from spHCN channels with L-Anap incorporated at different sites elicited by a series of hyperpolarizing voltage pulses from 0 mV to −120 mV, in the presence of 1 mM cAMP.

**Extended Data Fig. 3.**
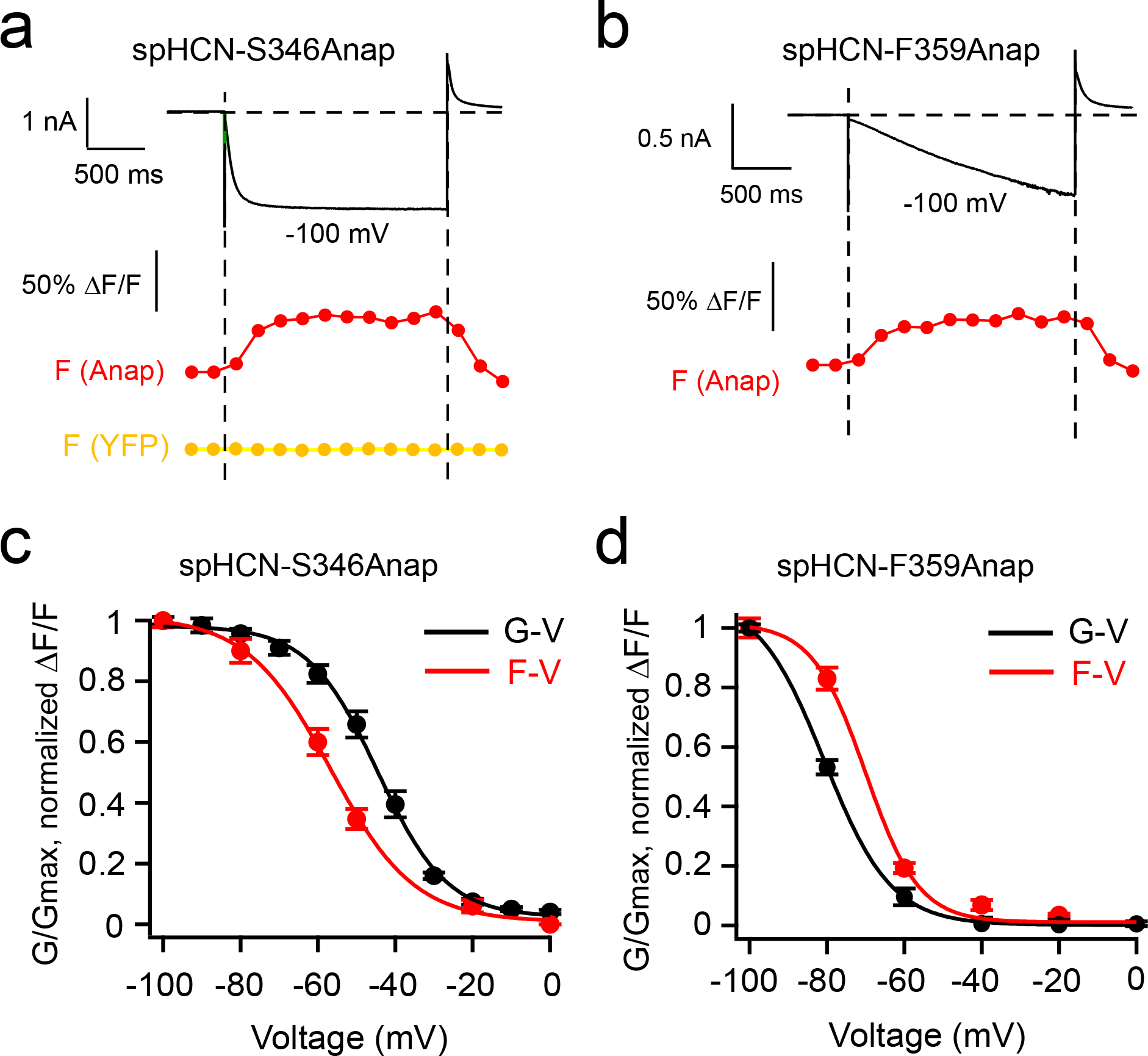
Hyperpolarization-dependence of the increase in Anap fluorescence. **a** and **b**, Simultaneous current and Anap fluorescence recordings of spHCN-S346Anap and spHCN-F359Anap channels in response to voltage steps to −100 mV, with 1 mM cAMP on the intracellular side. The YFP fluorescence from spHCN-S346Anap is also shown in the presence of cAMP. **c** and **d**, Fluorescence change-voltage (F-V) and conductance-voltage (G-V) relationships for the spHCN-S346Anap and spHCN-F359Anap channels, respectively. Because of its slow activation, the G-V curve for spHCN-F359Anap was not at steady state, therefore the F-V is shifted to more depolarized voltages compared to the G-V.

**Extended Data Fig. 4.**
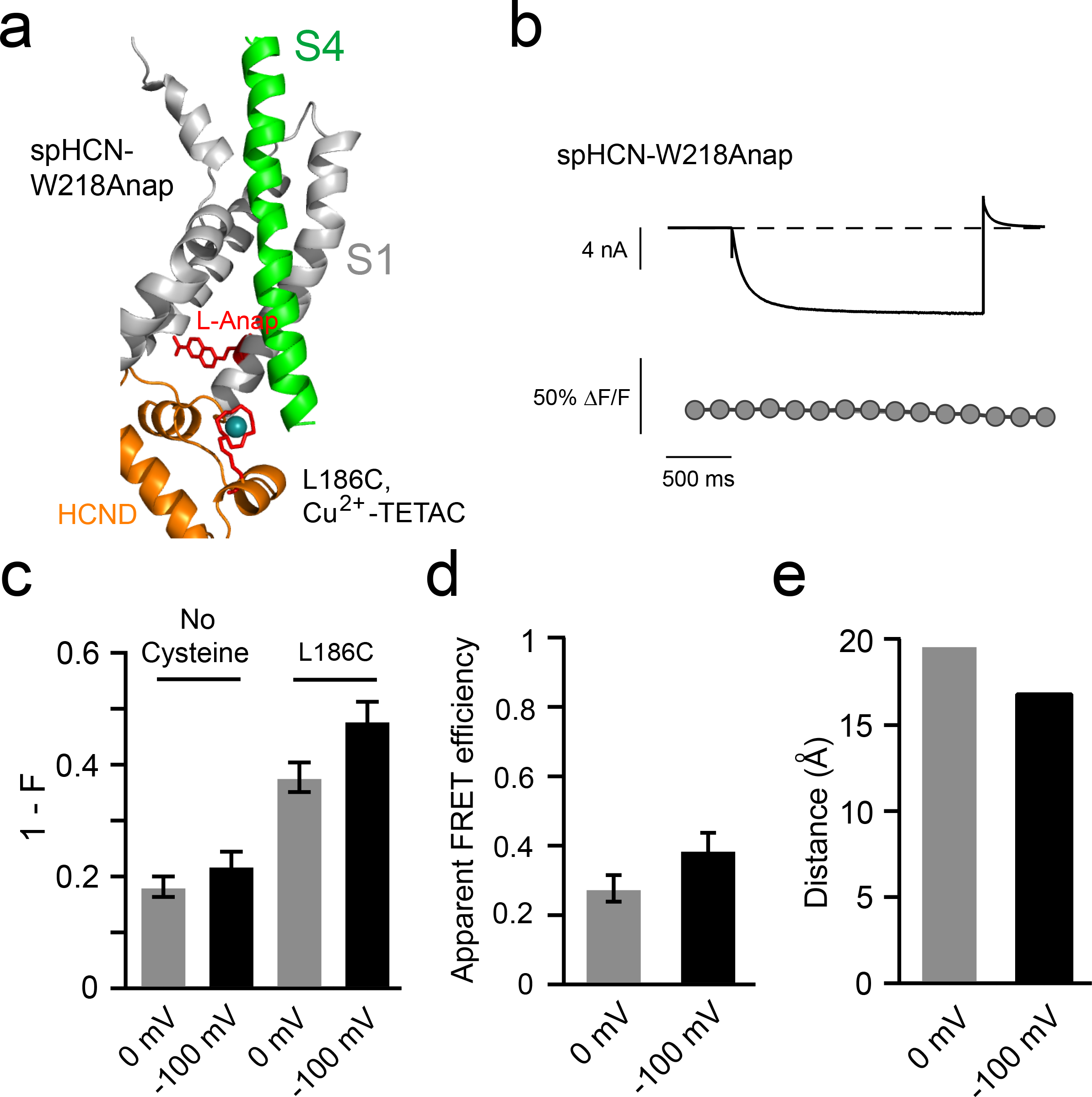
ACCuRET between an Anap site in the S1 helix and the transition metal ion site in the HCN domain. **a** Ribbon diagram showing the W218 Anap site and L186C modified by Cu^2+^-TETAC. **b** and **c**, Simultaneous current and Anap fluorescence recordings of spHCN-W218Anap channels. The Anap fluorescence did not change appreciably with −100 mV voltage pulses in the presence of 1 mM cAMP. **d**, Summary of the fractional Cu^2+^-TETAC quenching of Anap fluorescence at the W218 site, without and with the introduced cysteine L186C, at 0 mV and −100 mV (n = 4). **e**, FRET efficiency calculated using the quenching data in panel d and Equation 2 (see Methods). **f**, Summary showing the distances of the W218/L186 FRET pair at 0 mV and −100 mV, calculated from the FRET efficiency in panel e and the FCG plot in Fig. 3d.

**Extended Data Fig. 5.**
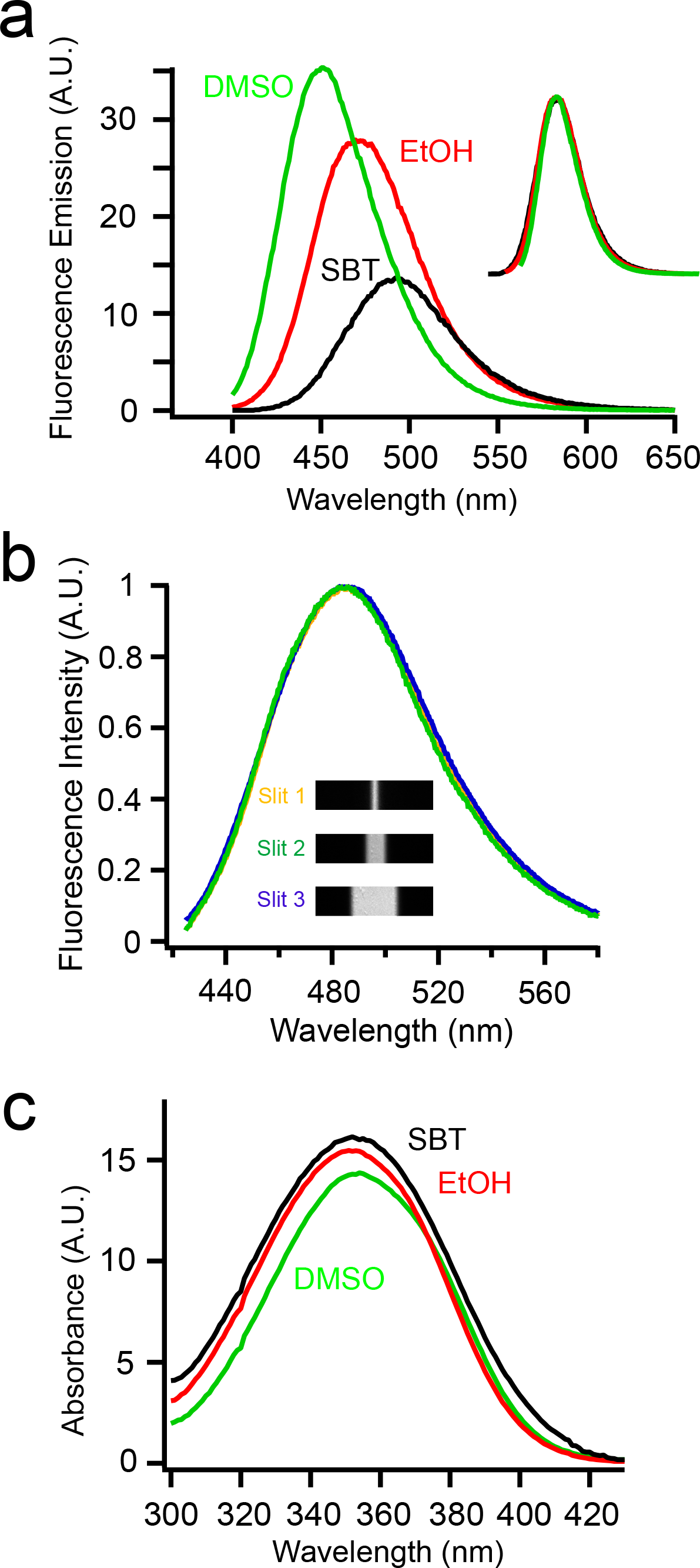
Fluorescence properties of L-Anap. **a**, Emission spectra of free L-Anap in different solvents. Insert: overlap of spectra in the different solvents illustrates that the shape of the spectra is not appreciably affected by different solvents. **b**, Emission spectra of L-Anap using the spectrograph attached to the patch-clamp microscope, using different slit widths. Slit 1 width: 5.9 μm; slit 2 width: 10.7 μm; slit 3 width: 24.8 μm when projected using a 60X water-immersed objective. Again, the shape of the spectra is not appreciably affected by different slit widths up to 25 μm, much larger than the diameter of our patches. **c**, Absorption spectra of free L-Anap in different solvents. In PCF experiments, we assume the change in the absorption of L-Anap incorporated into the channels is negligible compared to the environmentally-associated change in the quantum yield.

**Extended Data Fig. 6.**
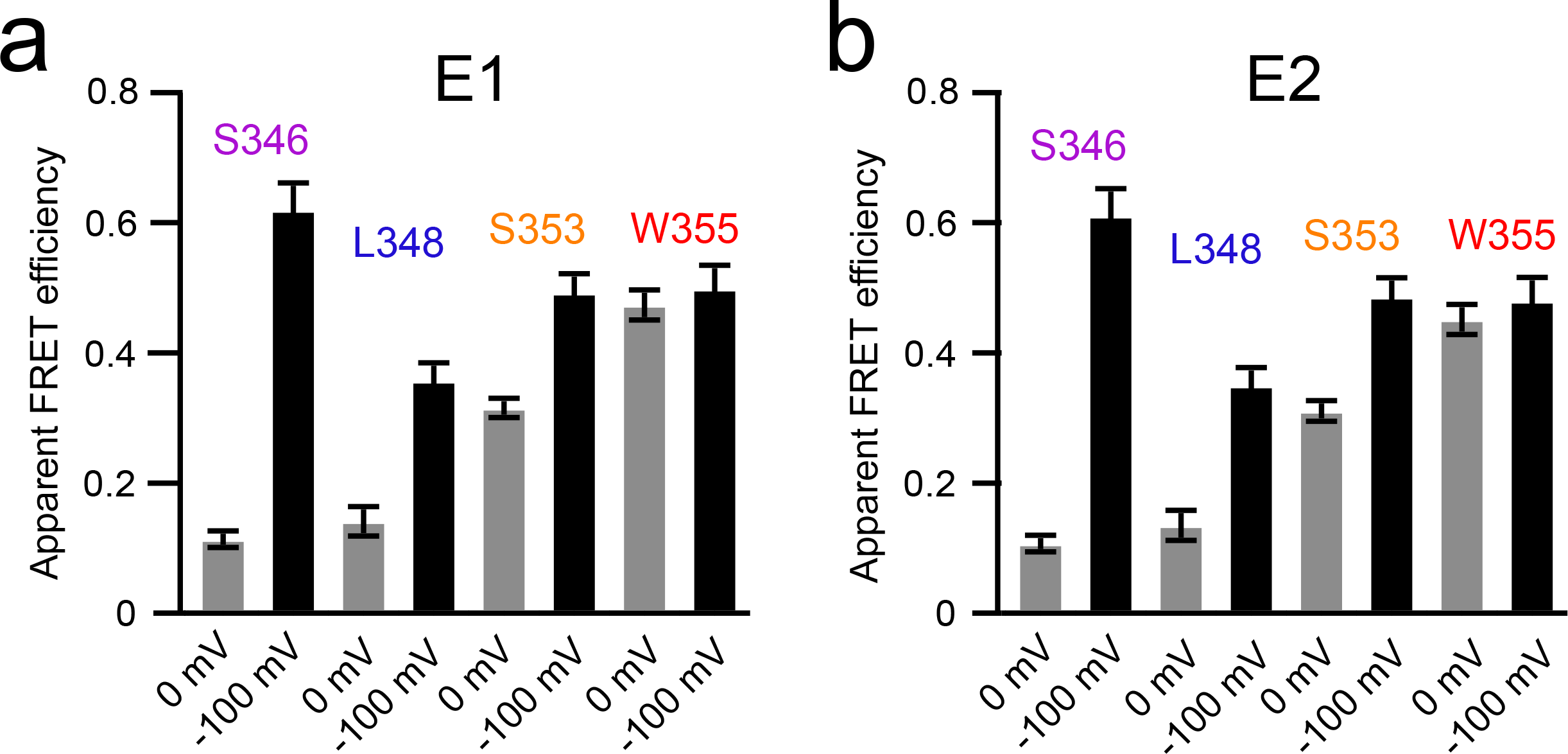
Different methods for correcting background quenching produce similar measurements of FRET efficiency. **a** and **b**, FRET efficiency of the various FRET pairs on spHCN channels based on the fractional quenching data in Fig 3c, calculated by the equations E1 and E2 (see Methods), respectively.

**Extended Data Fig. 7.**
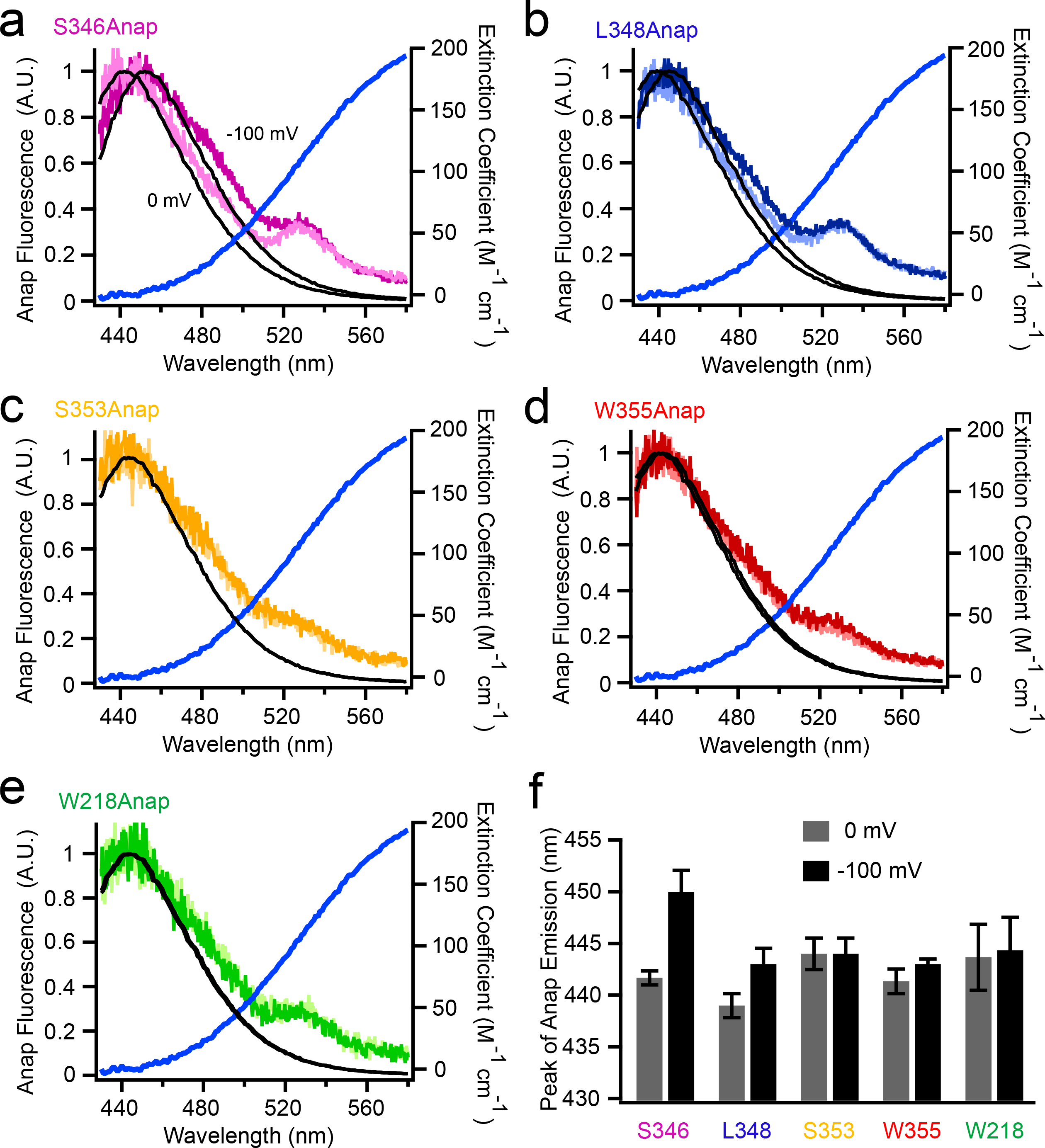
Emission spectra for L-Anap at each position in spHCN. **a-e**, Emission spectra of L-Anap measured with a spectrograph for the indicated positions in spHCN at 0 mV and −100 mV. These spectra were fit with emission spectra for free L-Anap in solution, measured with a fluorometer, and shifted so that the peak positions are the same (black traces). The small peak at about 530 nm most likely corresponds to direct excitation of YFP. Also shown is the absorption spectra of Cu^2+^-TETAC. **f**, Summary of the peak wavelength of the Anap emission for the various Anap sites in spHCN (n = 3-4).

**Extended Data Fig. 8.**
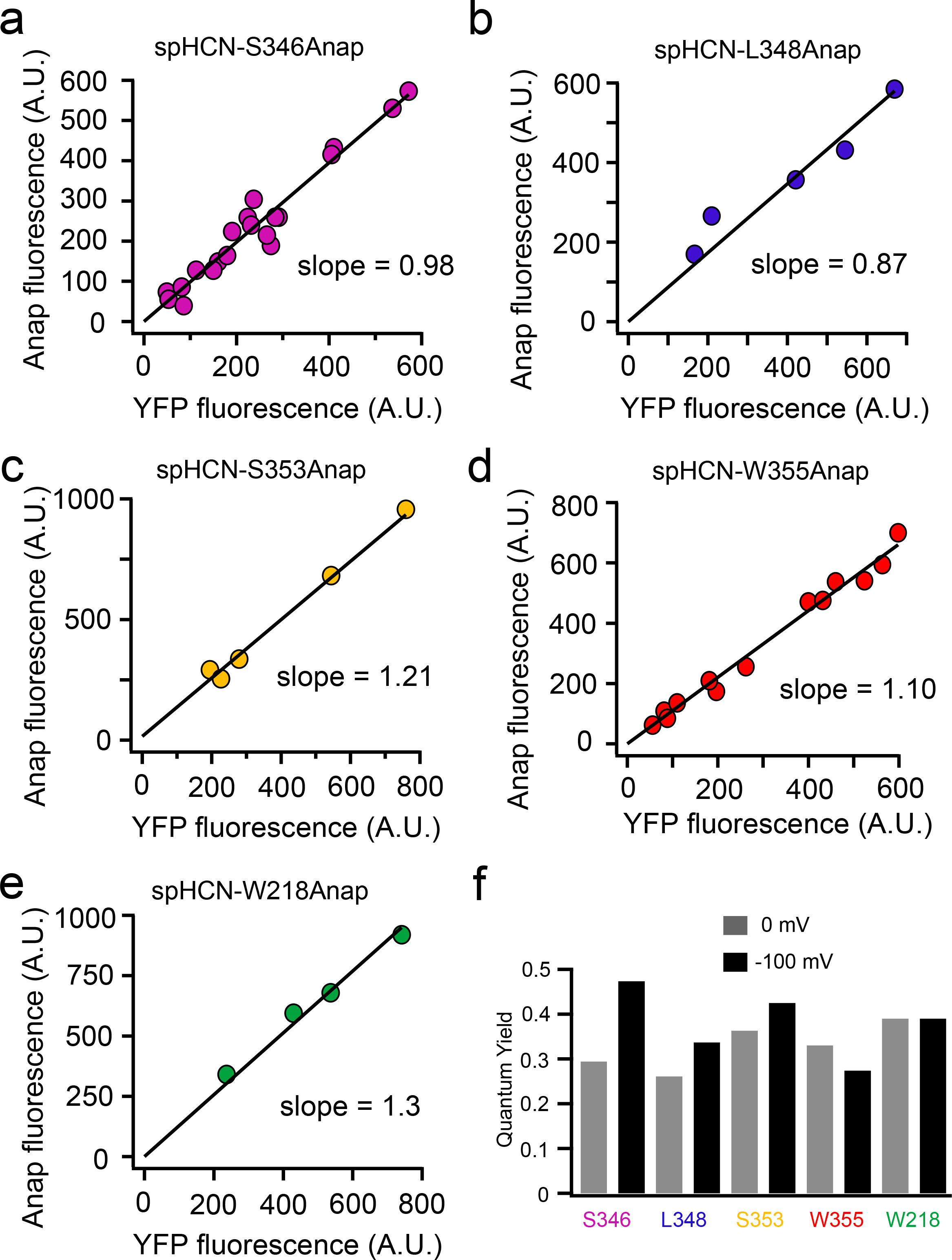
Correlation of the Anap fluorescence and the YFP fluorescence and estimation of the quantum yields of L-Anap. **a-e**, Plots of Anap versus YFP fluorescence intensities from multiple patches for the various L-Anap sites in spHCN. The slopes of the linear fit were used to calculate the L-Anap brightness relative to YFP. **f**, From the brightness of L-Anap at 0 mV and −100 mV, the quantum yields were estimated assuming the extinction coefficient of L-Anap was unchanged at different sites and different voltages.

## SUPPLEMENTARY VIDEO LEGENDS

### Video 1

Patch-clamp fluorometry image of spHCN-S346Anap channels in response to a repetitive −100 mV hyperpolarizing voltage in the presence of 1 mM cAMP in the bath. The upper left bar indicates three one-second pulses to −100 mV interspersed with one-second intervals at 0 mV. The holding potential is 0 mV. Scale bar on the lower left is 15 μm.

### Video 2

Patch-clamp fluorometry image of spHCN-W355Anap channels in response to a −100 mV hyperpolarizing voltage in the presence of 1 mM cAMP in the bath. The upper left bar indicates a two-second pulse to −100 mV (the same duration as in Fig. 1f and g). Scale bar on the lower left is 15 μm.

### Video 3

Rosetta model of the hyperpolarization-induced S4 voltage sensor movement in the spHCN channel. The movie shows a morph between the homology model based on the cryo-EM structure of hHCN1 (State 0) and the Rosetta model at −100 mV (State 2). The two models are shown in Fig. 4a.

### Video 4

The positions of the S4 voltage sensor from the spHCN channel in the experimentally-constrained 0 mV state (State 1) and the 0 mV state in the homology model based on the cryo-EM structure of hHCN1 (State 0). The two structures are quite similar with little translation of S4.

### Video 5

Interactions between arginines on S4 and aspartic acids within the S1-S3 helices during hyperpolarization-dependent activation of spHCN channels. The movie shows a morph between the 0 mV and −100 mV states, which are depicted in Fig. 4c.

### Video 6

Comparison of two different examples of S4 movement. Model of S4 movement in spHCN channels (left) and S4 movement in TPCs (right). The latter is a morph between the up state of VSD2 S4 in mouse TPC1 (PDB: 6C96) and the down state of the VSD2 S4 in Arabidopsis TPC1 (PDB: 5E1J).

